# Bacterial retrons function in anti-phage defense

**DOI:** 10.1101/2020.06.21.156273

**Authors:** Adi Millman, Aude Bernheim, Avigail Stokar-Avihail, Taya Fedorenko, Maya Voichek, Azita Leavitt, Rotem Sorek

## Abstract

Retrons are bacterial genetic elements comprised of a reverse transcriptase (RT) and a non-coding RNA. The RT uses the non-coding RNA as a template, generating a chimeric RNA/DNA molecule in which the RNA and DNA components are covalently linked. Although retrons were discovered three decades ago, their function remained unknown. In this study we report that retrons function as anti-phage defense systems. The defensive unit is composed of three components: the RT, the non-coding RNA, and an effector protein. Retron-containing systems are abundant in genomic “defense islands”, suggesting a role for most retrons in phage resistance. By cloning multiple retron systems into a retron-less *Escherichia coli* strain, we show that these systems confer defense against a broad range of phages, with different retrons defending against different phages. Focusing on a single retron, Ec48, we show evidence that it is a “guardian” of RecBCD, a complex with central anti-phage functions in the bacterial cell. Inhibition of RecBCD by dedicated phage proteins activates the retron, leading to abortive infection and cell death. Thus, the Ec48 retron forms a second line of defense that is triggered if the first lines of defense have collapsed. Our results expose a new family of anti-phage defense systems abundant in bacteria.

## Introduction

Retrons are genetic elements composed of a non-coding RNA (ncRNA) and a specialized reverse transcriptase (RT). These elements typically generate a chimeric RNA-DNA molecule, in which the RNA and DNA components are covalently attached by a 2′-5′ phosphodiester bond (Supplementary Figure 1). Retrons were originally discovered in 1984 in *Myxococcus xanthus*, when Inouye and colleagues identified a short, multi-copy single-stranded DNA (msDNA) that is abundantly present in the bacterial cell^1^. Further studies showed that this single stranded DNA is covalently linked to an RNA molecule^2^, and later deciphered in detail the biochemical steps leading to the formation of the RNA-DNA hybrid^3^. It was found that the retron ncRNA is the precursor of the hybrid molecule, and folds into a typical structure that is recognized by the RT^4^. The RT then reverse transcribes part of the ncRNA, starting from the 2’-end of a conserved guanosine residue found immediately after a double-stranded RNA structure within the ncRNA^3^. A portion of the ncRNA serves as a template for reverse transcription, which terminates at a defined position within the ncRNA^3^. During reverse transcription, cellular RNase H degrades the segment of the ncRNA that serves as template, but not other parts of the ncRNA, yielding the mature RNA-DNA hybrid^3^ (Supplementary Figure 1).

Dozens of retrons have been documented in a variety of microbial genomes, and 16 of them were studied experimentally in detail^5^. The documented retrons were all named following a naming convention that includes the first letters of their genus and species names, as well as the length of reverse-transcribed DNA (e.g., Ec48 is a retron found in *Escherichia coli* whose reverse transcribed DNA segment is 48 nt long). All studied retrons contain an RT and a ncRNA, with the conserved guanosine from which reverse transcription is initiated^6^. However, the sequences and lengths of the reverse transcribed template significantly vary and frequently show no sequence similarity between retrons^7^. The ability of retrons to produce ssDNA in situ has been adapted for multiple applications of synthetic biology and genome engineering^5,8–10^.

Although retrons have been studied for over 35 years, their biological function remained unknown. It has been suggested that retrons are a form of selfish genetic elements^11^, or have a function in coping with starvation^12^, pathogenesis^13^, and cell-specialization^5^. However, evidence for these functions were circumstantial and the mechanism by which retrons would exert these putative functions was not identified. In the current study we show that retrons form a functional component in a large family of anti-phage defense systems that are widespread in bacteria and confer resistance against a broad range of phages.

## Results

We initiated the current study by searching for reverse transcriptase genes that may participate in defense against phages. This search was inspired by prior reports on the involvement of RTs in bacterial defense^14–16^ and phage counter-defense mechanisms^17^. As bacterial defense systems tend to cluster in “defense islands” in microbial genomes^18–20^, we focused on RT genes of unknown function that are frequently encoded near known anti-phage systems such as restriction enzymes (Methods).

One of these RT genes is presented in Figure 1A-B. Homologs of this gene appear in a diverse set of bacteria, and show marked tendency to co-localize with known defense systems (Figure 1A). The RT gene is always found next to a second gene with a predicted ATPase domain, and we therefore hypothesized that the RT together with the ATPase form a two-gene phage resistance system. To test this hypothesis we cloned this two-gene system from *E. coli* 200499, together with its flanking intergenic regions, into the laboratory strain *E. coli* MG1655, which naturally lacks the system. We then challenged the transformed strain with a set of 12 coliphages and found that the system conferred protection against phages from a variety of families: T7 (*Podoviridae*), T4 and T6 (*Myoviridae*), and SECphi4, SECphi6, and SECphi18 (*Siphoviridae*) (Figure 1C, Supplementary Figure 2).

**Figure 1.**
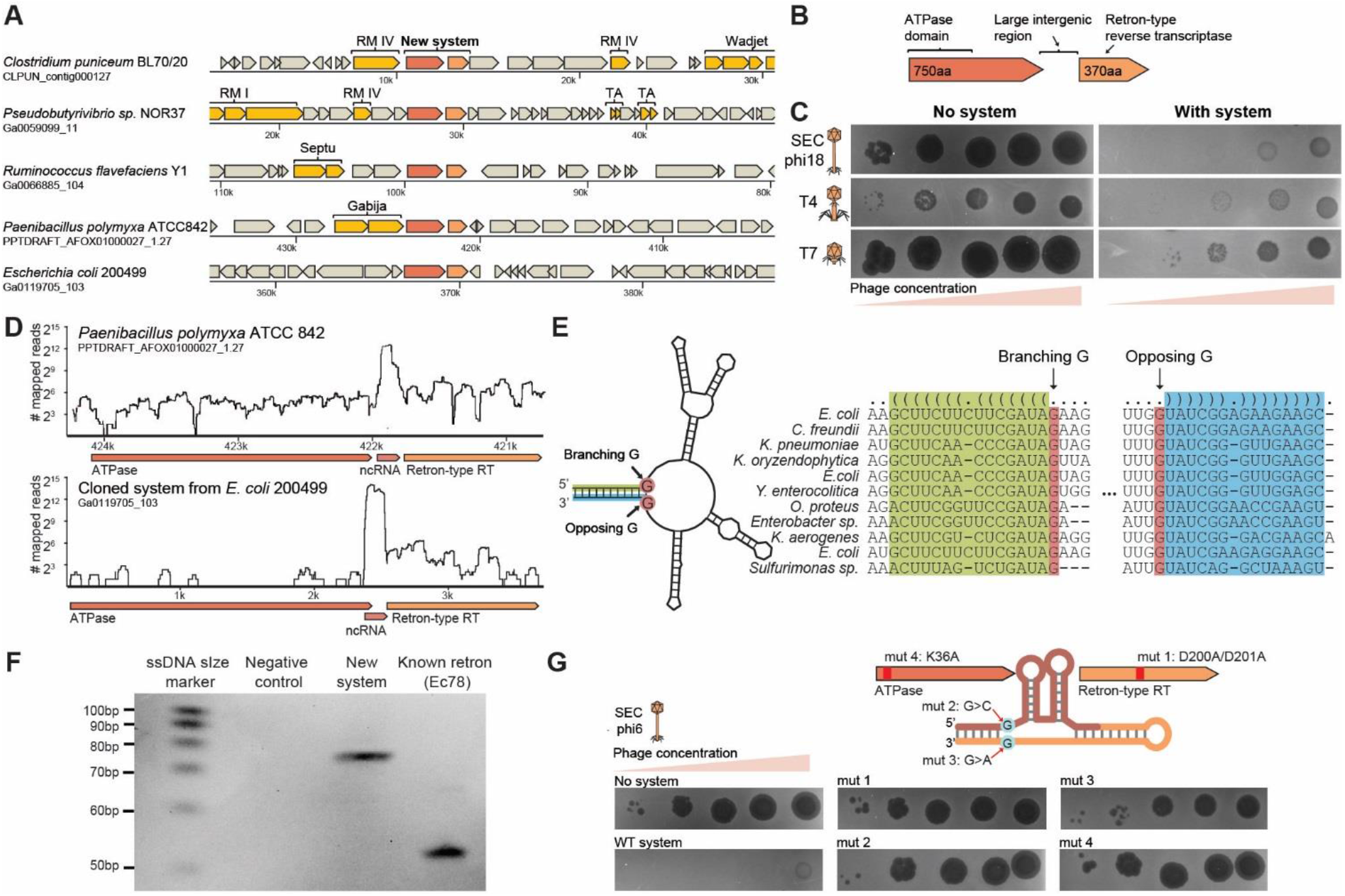
A retron-containing genetic system protects against phage infection. (A) A conserved two-gene cassette containing a reverse transcriptase is found in defense islands. Representative instances of the system, and their genomic environments, are presented. Genes known to be involved in defense are shown in yellow. RM, restriction modification; TA, toxin–antitoxin. Wadjet, Septu, and Gabija are recently described defense systems^19^. The encoding strain and the respective DNA scaffold accession on the IMG database^25^ are indicated to the left. (B) Domain organization in the gene cassette. (C) Serial dilution plaque assays shown for three phages on *E. coli* MG1655 strain transformed with the two-gene cassette (“with system”) or with a control plasmid encoding an RFP (“no system”). Images are representative of two replicates. (D) RNA-seq coverage of the systems loci in *Paenibacillus polymyxa* ATCC 842 (natively encoding the system, top) and *E. coli* MG1655 into which the system from *E. coli* 200499 was transformed (bottom). (E) Predicted structure of the ncRNA in the intergenic region of the tested system shows features of retron-type ncRNA (left). The structure corresponds to positions 369202-369366 in the *E. coli* 200499 genome assembly (scaffold Ga0119705_103). Alignment and structure prediction of the intergenic region from homologs in multiple bacteria are shown on the right. Areas of conserved base pairing are highlighted. Predicted structure is presented on the top, with each pair of parentheses representing base-pairing. (F) Production of multicopy single stranded DNA (msDNA). The msDNA was extracted from *E. coli* strains expressing the ncRNA and RT of the new defense system, a known retron Ec78, or a negative control expressing GFP instead (Methods). Extracted msDNA was examined on a 10% denaturing polyacrylamide gel. (G) Mutational analysis of elements within the new defense system. Shown are serial dilution plaque assays with phage SECphi6, comparing the strain with the wild type system to strains with mutated versions. Images are representative of two replicates.

The presence of an atypically large (>100 nt), conserved intergenic region between the ATPase and RT genes led us to hypothesize that this intergenic region might contain a non-coding RNA (Figure 1B). Indeed, examining RNA-seq data from *Paenibacillus polymyxa*^21^, which naturally encodes this system, showed high levels of expression from the intergenic region (Figure 1D). Similar expression patterns were observed in RNA-seq data from the *E. coli* strain into which we cloned the defense system, consistent with the presence of a ncRNA in the intergenic region (Figure 1D).

Because the RT gene showed significant homology to retron-type RTs^22^, we hypothesized that the newly discovered defense system contains a retron, and that the ncRNA we detected is the retron ncRNA precursor. In support of this, we found that the predicted secondary structure of the ncRNA conforms with the characteristics of known retron ncRNA precursors, including the conserved local dsRNA structure immediately followed by non-paired guanosine residues on both strands (Figure 1E; Supplementary Figure 1). These structural features were conserved among homologs of this system (Figure 1E). To check if the ncRNA indeed forms a precursor for single-stranded DNA (ssDNA) synthesis, we extracted ssDNA from a strain into which we cloned the RT and ncRNA, and found a ssDNA species sized between 70-80 nt, which was absent from the control strain that contained a GFP gene instead (Figure 1F). These results confirm that the new defense system we discovered contains a previously unidentified retron.

To examine whether the retron features are involved in the anti-phage activity of the new defense system, we experimented with mutated versions of the system. Point mutations in the conserved YADD motif of the catalytic core of the RT (D200A and D201A)^23^, rendered the system inactive (Figure 1G). Similarly, a point mutation in the ncRNA, mutating the guanosine predicted as the branching residue priming the reverse transcription^24^ (G>C at position 17 of the ncRNA), or the second conserved guanosine that was shown in other retrons to be essential for initiation of reverse transcription^24^ (G>A at position 147), completely abolished defense against phages (Figure 1G). These results suggest that proper reverse transcription of the retron ncRNA is essential for its defensive function. We also found that a point mutation in the ATP-binding motif of the associated ATPase gene (K36A) completely eliminated the defense phenotype, showing that the ATPase gene is an indispensable component of the retron-containing defense system. We therefore conclude that the new defense system consists of three components essential for its anti-phage activity: the RT and ncRNA (which together form an active retron) and an additional gene that contains an ATPase domain. Following a recently proposed revised nomenclature for retrons^5^ we termed this defense system Retron-Eco8.

The identification of a novel defense system that contains a retron led us to ask whether retrons in general may have a role in defense against phages. If this is the case, we would expect to find retrons enriched in defense islands, near known anti-phage defense systems. To test this hypothesis we searched for homologs of the RT proteins of previously characterized retrons, in a set of 38,167 bacterial and archaeal genomes (Methods). We detected 4,802 homologs of retron RTs in 4,446 of the genomes, and used hierarchical clustering to divide these homologs into eight clades (Figure 2A, Supplementary Table 1). We found that in six of the clades, the RT genes had a strong tendency to be genomically associated with other known defense genes in defense islands (Methods). Between 38% and 47% of the genes in each of these clades were found to be located near known defense systems, implying that most retrons may participate in anti-phage defense.

**Figure 2.**
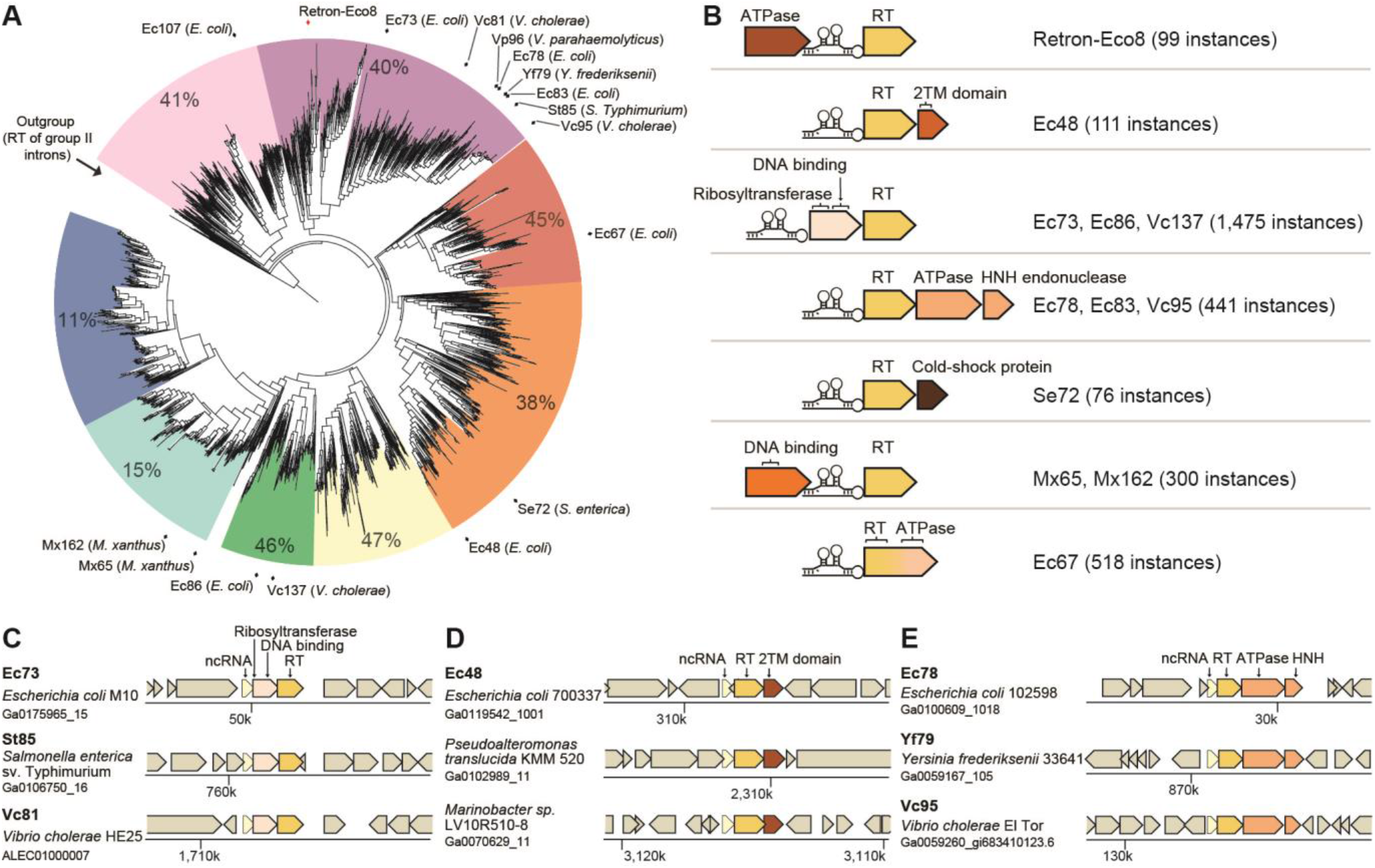
Known retrons are part of multi-gene systems and are located in defense islands. (A) Phylogenetic analysis of homologs of retron RTs. Known retrons are marked around the tree with the encoding species stated in parentheses. For each clade, the percentage of genes that appear next to known defense systems is presented (see Methods). (B) Types of retron-containing systems, classified based on the identity of the associated gene. The number of identified RT homologs with each gene configuration is presented in parentheses. (C-E) The genomic environments of known retrons and their homologs, divided into systems containing an associated ribosyltransferase (C), a two-transmembrane (2TM) domain gene (D), or genes with ATPase and HNH endonuclease domains (E). Names of known retrons are in bold. The encoding strain and the respective DNA scaffold accession in the IMG database^25^ are indicated for each retron or retron homolog.

Retrons have been previously described as two-component systems, comprised of the RT and the precursor ncRNA^5^. However, when examining the genomic environments of known retrons and their homologs, we observed that the vast majority are encoded as part of a gene cassette that includes one or two additional protein-coding genes (Figure 2B-E). For example, retrons Ec73, St85, and Vc81 (found in *E. coli*, *Salmonella enterica*, and *Vibrio cholerae*, respectively) all have an upstream gene containing a ribosyltransferase and DNA-binding domains (Figure 2C); and retron Ec48 and its homologs are linked to a gene encoding a protein with two predicted transmembrane (2TM) helices (Figure 2D). Some retrons, including Ec78, Yf79, and Vc95 are encoded as part of a cassette that includes two additional genes: one with an ATPase domain and another with an HNH endonuclease domain, a gene organization that was previously identified in the anti-phage system Septu^19^ (Figure 2E). In other cases, for example retron Ec67, the associated gene is fused to the RT gene (Figure 2B). The tight genetic linkage of retrons with these genes suggests that the functional unit that includes the retrons also includes the associated genes. We refer to these associated genes as retron “effectors”, due to reasons explained below.

To test whether retrons function as anti-phage defense systems, we experimentally examined the previously characterized retrons that were identified in *E. coli* strains (six retrons, excluding Ec107, see Methods) as well as a number of validated retrons encoded by *S. enterica* and *V. cholerae* (five additional retrons). We cloned each retron, together with the predicted effector gene(s), into an *E. coli* MG1655 strain that is not known to encode retrons. We then challenged the retron-containing bacteria with an array of 12 phages that span several major phage families. For eight of the 11 systems, we observed marked anti-phage activity against at least one phage (Figure 3, Supplementary Figure 3). For some strains (e.g., those harboring retrons Ec86 and Se72), anti-phage activity was restricted to one phage, while for others (Ec73 and Ec48) defense was broad, spanning phages from several different phage families (Figure 3, Supplementary Figure 3).

**Figure 3.**
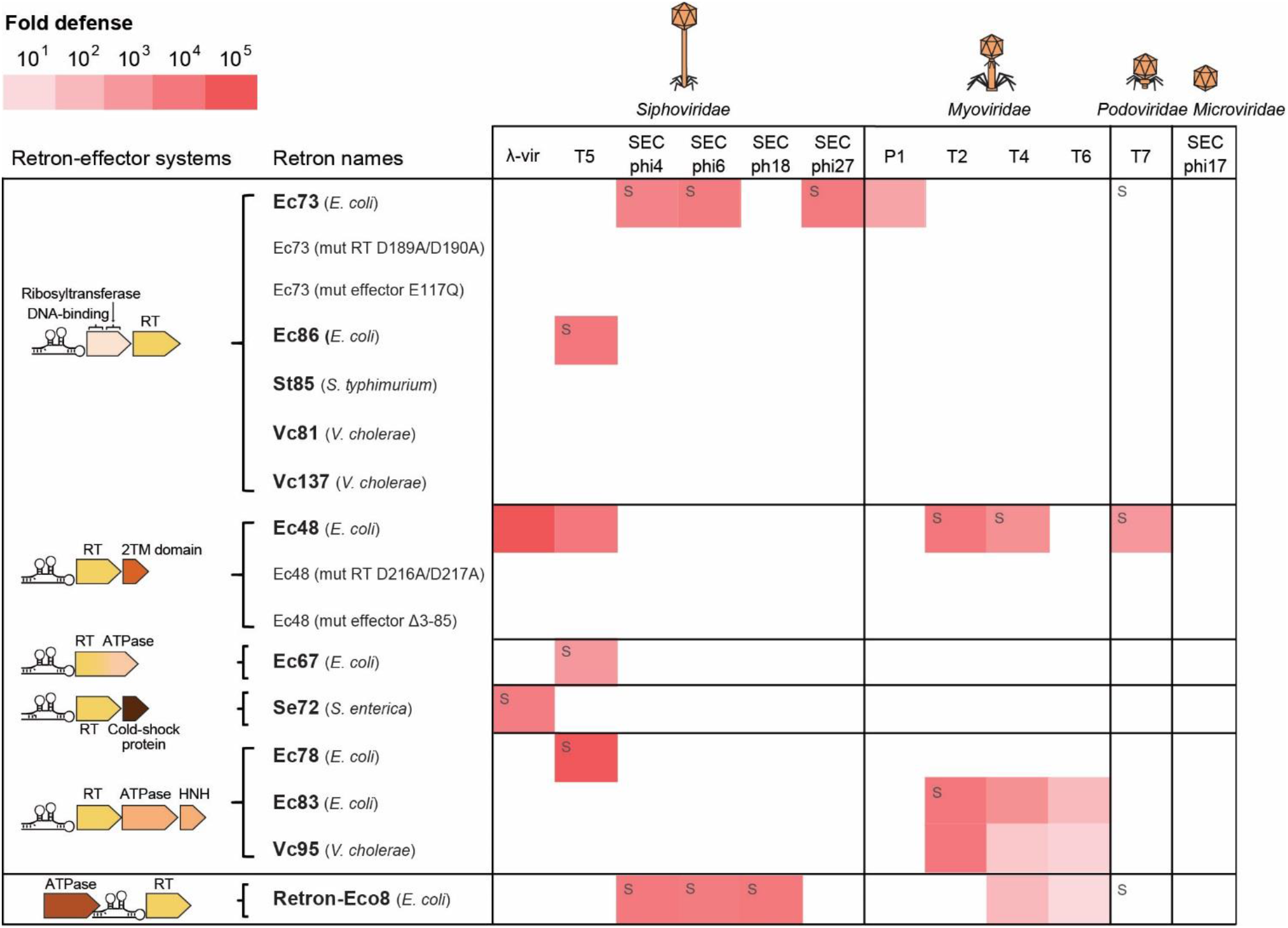
Known retrons, together with their effector genes, protect against phages. Fold protection was measured using serial dilution plaque assays, comparing the efficiency of plating (EOP) of phages on the system-containing strain to the EOP on a control strain that lacks the system. Data represent an average of two replicates (Supplementary Figure 3). A designation of ‘s’ stands for a marked reduction in plaque size. On the left, gene organization of the defense systems, with identified domains indicated. For retrons Ec73 and Ec48, data for WT strains are presented as well as data for strains mutated in either the reverse transcriptase (RT) or the associated gene (effector). The Δ3-85 mutation in the effector gene of Ec48 reflects a deletion of the two predicted transmembrane helices.

To assess whether both the retron activity and the activity of its effector gene are necessary for anti-phage defense, we further experimented with the two retron systems that showed the broadest defense (Ec48 and Ec73). Point mutations predicted to inactivate the catalytic site of the RT (D216A/D217A in the RT of Ec48, and D189A/D190A in the RT of Ec73) completely abolished defense, indicating that reverse transcription of the retron is essential for defense. In addition, an E117Q point mutation predicted to inactivate the catalytic site of the ribosyltransferase domain of the Ec73 effector^26^, led to a non-functional system, and similarly, deletion of the transmembrane helices of the gene associated with the Ec48 retron also abolished defense (Figure 3, Supplementary Figure 3). Together, these results show that retrons, functioning together with their associated effector genes, form anti-phage defense systems.

To gain further insight into the mechanism by which retrons recognize and mitigate phage infection, we attempted to find phage mutants that escape retron defense. For this, we focused on retron Ec48, as it provided strong defense against phages belonging to three different families (*Siphoviridae*, *Myoviridae* and *Podoviridae)* (Figure 3). We were able to isolate six mutants of phage λ-vir, as well as two mutants of phage T7, which could overcome the defense conferred by Ec48 and its effector gene (Figure 4A). We then sequenced the full genome of each of the phage mutants, and compared the resulting sequences to the sequence of the wild-type (WT) phage. In five of the six retron-overcoming mutants of phage λ-vir, we found a single point mutation that distinguished the mutant phage from the WT phage. In all five cases, the point mutation (either a single base deletion or a single base insertion) resulted in a frame shift in a λ-vir gene called *gam*; and in the sixth mutant we detected 16 mutations, one of which also causing a frameshift in *gam* (Figure 4A, Supplementary Table 2). These results suggest that mutations that inactivate the Gam protein of phage λ-vir enable the phage to overcome the defensive activity of the retron Ec48 defense system. In the retron-escaping T7 mutants we found two missense point mutations appearing in both mutants, one in gene *1.7* (tyrosine at position 128 mutated into cysteine) and the other in gene *5.9* (changing a leucine in position 23 of the protein to proline) (Figure 4A, Supplementary Table 2).

**Figure 4.**
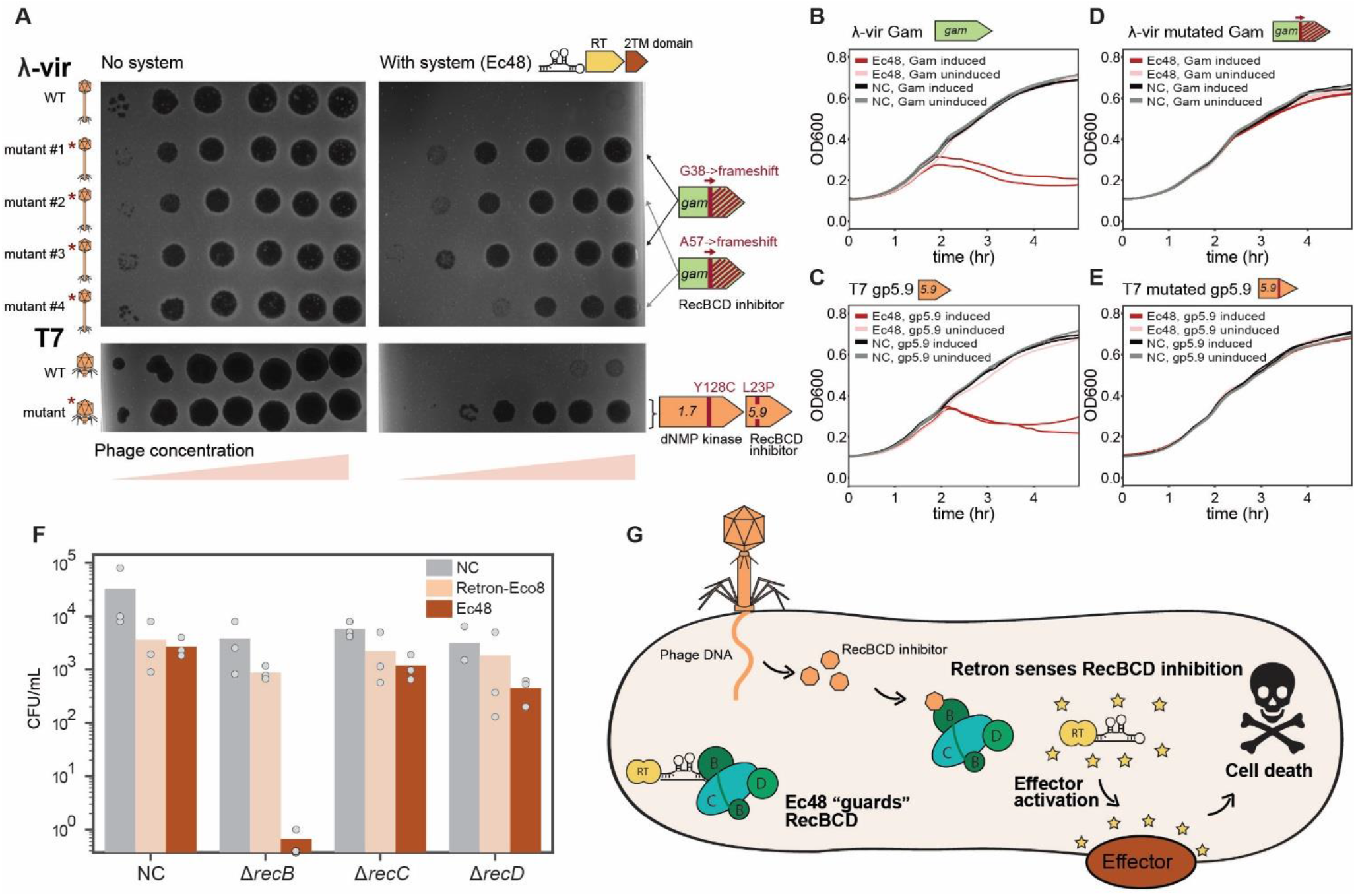
The Ec48 retron anti-phage system functions through abortive infection. (A) Representative phage mutants capable of escaping Ec48 retron system defense. Shown are serial dilution plaque assays, comparing the plating efficiency of WT and mutant phages on bacteria that contain the Ec48-retron anti-phage system and a control strain that lacks the system and contains an empty vector instead. Images are representative of three replicates. Genes mutated in mutant phages are presented on the right. λ-vir mutant #1 contains 15 additional mutations as detailed in Supplementary Table 2. (B-E) Expression of phage-encoded RecBCD inhibitors activates cell death in Ec48-containing strains. Growth curves of *E. coli* expressing the Ec48 system or a negative control vector that lacks the system (NC). The expression of Gam protein of phage λ (B), gp5.9 protein of phage T7 (C), or their mutated versions (panels D-E) was induced by the addition of 0.3% arabinose to exponentially growing cells at OD_600_ 0.3. For each experiment, two biological replicates are presented as individual curves, each a mean of two technical replicates. (F) A vector that contains the Ec48 anti-phage system cannot be transformed into cells lacking RecB. Transformation efficiency of a plasmid containing either the Ec48 retron system, the newly discovered anti-phage retron (Retron-Eco8) or RFP (as a negative control, NC), into *E. coli* strains with a deletion in one of the RecBCD components or a deletion of iscR (an unrelated bacterial gene as a negative control, NC). The number of colony forming units following transformation is presented. Bar graph represents an average of three replicates, with individual data points overlaid. (G) Model for the anti-phage activity of the Ec48 retron system.

Remarkably, both the Gam protein of phage λ and the gp5.9 protein of phage T7 have the same biological role: inhibition of a bacterial complex called RecBCD^27–29^. The RecBCD complex is known to have a central role in DNA repair as well as in anti-phage activity^30,31^. RecBCD rapidly degrades linear dsDNA and hence phage DNA, which is usually injected into the infected cell in a linear form, is susceptible to RecBCD-mediated degradation^30^. RecBCD also plays an important role in acquisition of new CRISPR spacers from phage genomes^32^ and in the formation of guide DNAs for prokaryotic argonaute proteins that defend against foreign nucleic acids^33^. Since RecBCD is a central immunity hub in the bacterial cell, numerous phages are known to encode anti-RecBCD proteins (including the λ Gam and T7 gp5.9 proteins), which bind the RecBCD complex and inactivate it^30,34^. Our findings that mutations in RecBCD inhibitors of both phage λ-vir and phage T7 enabled the phages to escape defense suggest that the Ec48 retron senses these phage proteins, or their activities, as a signal for phage infection.

The Ec48 retron defense system contains a putative effector gene with two transmembrane-spanning helices (Figure 2D). Such a transmembrane-spanning domain organization is common in effector proteins of CBASS anti-phage defense systems that cause the cell to commit suicide once phage infection has been sensed^20^. In CBASS, the transmembrane-spanning protein is responsible for the cell-killing effect, and is predicted to form pores in the membrane after receiving a signal indicative of phage infection, causing the infected bacteria to die before the phage is able to complete its replication cycle^20^. We therefore hypothesized that the Ec48 retron defense system also functions by inflicting cell death, and that the cell-suicide transmembrane-spanning effector is activated when the retron senses the phage RecBCD inhibitors.

To test this hypothesis, we expressed the RecBCD inhibitors from phages λ-vir and T7 (Gam and gp5.9, respectively) in cells that contain the Ec48 defense system, and followed bacterial growth after induction of expression. We observed a marked decrease in bacterial cell density following the expression of either of the two RecBCD inhibitors in bacteria that contain the Ec48 retron defense system, but not in bacteria lacking Ec48 (Figures 4B-C). Expression of the mutated versions of the phage RecBCD inhibitor genes, which enabled them to escape from Ec48 defense, did not activate bacterial suicide (Figure 4D-E). To verify that the observed reduction in the bacterial cell density corresponds to cell death, we induced the expression of the T7 RecBCD inhibitor gp5.9 in bacteria that contain the Ec48 defense system, and plated bacteria to count colony forming units (CFUs). Counts of viable bacteria dropped dramatically following induction of T7 gp5.9 in bacteria that contained the retron system, consistent with cell death (Supplementary Figure 4A). Altogether, these results indicate that once the Ec48 retron defense system senses the presence of phage-encoded RecBCD inhibitors, it causes cell death. This form of protection against phage infection is called abortive infection (Abi), and is a common defensive strategy employed by multiple, mechanistically diverse bacterial immune systems^35^.

Notably gp1.7, the second gene found to be mutated in the Ec48-escaping T7 phages, did not cause cell death when expressed in Ec48-containing cells (Supplementary Figure 4B). Perhaps this mutation allows the phage to escape a different step in the Ec48 defense process, or, alternatively, this mutation may be a passenger mutation not involved in escape from retron defense.

Although the Gam protein of phage λ and the gp5.9 protein of phage T7 both activate the Ec48 retron defense system to induce cell death, these proteins share no detectable sequence similarity. We therefore hypothesized that rather than directly sensing the phage proteins, the Ec48-containing system monitors the integrity of RecBCD itself, and becomes activated when RecBCD is tampered with. If this hypothesis is correct, then perturbations of RecBCD that are not linked to phage infection would also trigger the Ec48 defense system and lead to cell death. To test this hypothesis we used strains of *E. coli* in which components of RecBCD are deleted. Such cells are impaired in DNA repair but are viable in non-stress conditions^36,37^. We attempted to transform a plasmid that encodes the Ec48 defense system or a control plasmid (lacking the system) into *E. coli* strains deleted for one of the three components of RecBCD (either Δ*recB*, Δ*recC* or Δ*recD*) or a control strain. While the control plasmid was transformed with high efficiency into all strains, transformation of the Ec48-containing plasmid into Δ*recB* consistently and repeatedly failed, yielding zero colonies or a single colony in three independent transformation attempts (Figure 4F). The same Ec48-containing plasmid was transformed with high efficiency into the Δ*recC*, Δ*recD* and the control strains (Figure 4F). These results suggest that the Ec48 system becomes activated specifically when the RecB protein is missing or impaired. Notably, the Gam protein of phage λ has been shown to inhibit RecBCD by directly binding to RecB^38^, supporting this model.

Combined together, our results lead to a mechanistic model in which the Ec48 retron defense system “guards” the normal activity of the bacterial immunity hub RecBCD. Inactivation of the RecB protein by phage inhibitors triggers the retron system and leads to cell suicide before the phage is able to complete its replication cycle (Figure 4G). As the cell-killing function is likely carried out by the effector transmembrane-spanning protein, we further hypothesize that the role of the reverse-transcribed retron RNA, perhaps together with the RT, is to sense or monitor the integrity of RecB. Notably, the RecB protein has a domain that binds ssDNA, to which the Gam protein has been shown to bind. Perhaps the reverse transcribed ssDNA in the retron binds RecB, and during infection it is displaced by the phage Gam protein, releasing the retron to somehow activate the effector protein.

Interestingly, a plasmid encoding the new retron defense system Retron-Eco8 was readily transformed into all three RecBCD deletion strains, including Δ*recB*, suggesting that the Retron-Eco8 defense system does not guard RecBCD and likely senses phage infection in a different manner (Figure 4F). Therefore, retron-containing defense systems other than Ec48 may sense alternative signals or guard other central components of the cell to mitigate phage infection.

## Discussion

Our results show that retrons broadly function in anti-phage defense, solving a three-decade-old mystery on the function of these peculiar genetic elements. The functional retron anti-phage system includes three components: the RT, the ncRNA, and an effector protein. Accessory open reading frames have recently been predicted as genetically linked to some retrons and were suggested to assist the retron function^5^; our data indicate that these accessory genes function as effector proteins essential for the anti-phage activity.

Interestingly, retron systems that have a similar genetic composition defend against different phages. For example, the Ec73 and Ec86 systems, both of which contain a ribosyltransferase effector, protect against a completely different set of phages (Figure 3, Supplementary Figure 3). The fact that these systems both encode genes with a similar effector domain, but encode retron ncRNA sequences that are significantly different and cannot be aligned, suggests that the nucleic acid component of the retron defense system and/or the RT participate in the recognition of phage infection (Figure 4G). The exact mechanism by which the retron RNA-DNA hybrid takes part in this recognition, and the mechanism by which the signal is transferred to the effector protein to induce cell death, remains to be elucidated. It is possible that the retron ssDNA component attaches to DNA-binding cellular complexes (such as RecBCD in Ec48) or phage proteins (in other retrons) to monitor their activities. It is also possible that the msDNA or some fragments of it are released during infection, and bind the effector protein to induce its activity. In support of this hypothesis, some of the effector proteins encode predicted nucleic acid binding domains (Figure 2B).

Our systematic search for homologs of retron RTs retrieved 4,802 homologs in more than 4,000 bacterial genomes that belong to many different phyla (Supplementary Table 1), suggesting that retrons form a widespread novel class of anti-phage defense systems (Figure 2A). In addition to the effectors identified next to experimentally verified retrons (Figure 2B), additional types of effectors can be seen next to homologs of retron RTs that were not yet experimentally studied. These include Toll/interleukin-1 receptor (TIR) and protease domains, both are known to be present in CBASS effector genes^20^ as well as in other phage-defense systems^19^. Therefore, retron defense systems present a variety of effector proteins that are predicted to exert the cell-suicide function of the system.

While most of the retron systems we tested provided defense against phages (Figure 3) and many retrons tend to appear in defense islands, two clades of RT homologs do not follow this pattern and are only rarely found next to known defense systems (Figure 2A). This suggests that some retrons may have adapted to perform functions other than anti-phage defense, perhaps in a manner analogous to toxin-antitoxin systems, some of which are involved in anti-phage defense while others play a role in the bacterial stress response^39^.

Our results suggest that Ec48 functions as a “guardian” of RecBCD, a central cellular hub of anti-phage immunity in *E. coli*. Such a defensive strategy makes immunological sense, because different phages can inhibit RecBCD by multiple different mechanisms^30,34^. Instead of recognizing the phage inhibitors themselves, which can be highly divergent and share no similarity, Ec48 monitors the integrity of the bacterial immune system, and if the integrity is impaired the molecular “conclusion” is that the cell has been infected by phage. An analogous strategy is exerted by the PrrC protein in *E. coli*, which similarly guards the normal function of type I restriction-modification systems and inflicts cell-suicide when phage proteins inhibit the restriction enzyme^40^. Remarkably, such a defensive strategy has been documented as a central aspect of the immune system of plants, where it was termed “the guard hypothesis”^41^. In plants, pathogen resistance proteins are known to guard central cellular processes and detect their disruption by pathogen effectors as a signature for infection^42^. A similar strategy has also been recognized in components of the animal innate immune systems, where the recognition of a pathogenicity signature often leads to inflammatory cell death^42^. Our discovery therefore shows that similar immunological principles govern the design of immune systems across bacteria, plants, and animals.

## Acknowledgements

We thank the Sorek laboratory members for comments on earlier versions of this manuscript. We also thank Hyeim Jung for providing us with a protocol for msDNA isolation. A.M. was supported by a fellowship from the Ariane de Rothschild Women Doctoral Program and, in part, by the Israeli Council for Higher Education via the Weizmann Data Science Research Center. A.B. is the recipient of a European Molecular Biology Organization (EMBO) Long Term Fellowship (EMBO ALTF 186-2018). R.S. was supported, in part, by the Israel Science Foundation (personal grant 1360/16), the European Research Council (grant ERC-CoG 681203), the Ernest and Bonnie Beutler Research Program of Excellence in Genomic Medicine, the Minerva Foundation with funding from the Federal German Ministry for Education and Research, and the Knell Family Center for Microbiology.

## Material and Methods

### Detection of RT genes in defense islands

Protein sequences of all genes in 38,167 bacterial and archaeal genomes were downloaded from the Integrated Microbial Genomes (IMG) database^25^ in October 2017. These proteins were filtered for redundancy using the ‘clusthash’ option of MMseqs2 (release 2-1c7a89)^43^ using the ‘–min-seq-id 0.9’ parameter and then clustered using the ‘cluster’ option, with default parameters. Each cluster with >10 genes was annotated with the most common pfam, COG, and product annotations in the cluster. For each cluster annotated as reverse transcriptase a defense score was calculated as previously described^19^, recording the fraction of genes in each cluster that have known defense genes in their genomic environment spanning 10 genes upstream and downstream the inspected gene. Members of clusters with high tendency to be associated with known defense genes were scanned for conserved gene cassettes as previously described^19^.

### Prediction of ncRNA structure

The upstream boundaries of Retron-Eco8 ncRNA were estimated based on a promoter prediction upstream to the RT of *E. coli* 200499 using BPROM^44^. The start position was chosen 10nt downstream to the predicted −10 sequence. The last position in the intergenic region before the RT gene was taken as the end point. The resulting predicted ncRNA corresponds to coordinates 369202-369366 (positive strand) in the *E. coli* 200499 assembly, scaffold accession number Ga0119705_103 in the IMG database^25^.

To predict the structure of the Retron-Eco8 ncRNA the extracted sequence was folded using the RNAfold web server^45^. For the alignment and structure prediction appearing in Figure 1E, the 250nt upstream of the RTs of homologs of the system were extracted. The sequences were filtered for redundancy using blastclust (with options −S 100 −L 1). The remaining sequences were analyzed using WAR web server^46^ and the MAFFT+RNAalifold representative results are shown.

### Genomic identification and analysis of retron RT homologs

A list of RTs of experimentally verified, known retrons (Mx162, Sa163, Mx65, EC67, EC48, EC86, Vc137, EC73, Vc81, St85, Ec78, EC83, YF79, Vc95, Vp96, EC107, Se72) together with the RT of Retron-Eco8 were searched against the downloaded database of protein sequences from 38,167 bacterial and archaeal genomes using the ‘search’ option of MMseqs2 (release 6-f5a1c). Hits were mapped to the clusters mentioned above, and protein clusters with at least 10 hits were taken as homologs. The genomic environments spanning 5 genes upstream and downstream of each of the homologs were searched to identify conserved gene cassettes.

To generate the phylogenetic tree in Figure 2, sequences shorter than 200aa were removed. The ‘clusthash’ option of MMseqs2 (release 6-f5a1c) was then used to remove protein redundancies (using the ‘–min-seq-id 0.9’ parameter). Six sequences of group II intron reverse transcriptase (WP_138709943, WP_137561357, WP_131190414, WP_130644660, WP_130706420, WP_130793790) were added and were used as outgroup. Sequences were aligned using MUSCLE (v3.8.1551)^47^ with default parameters. The FastTree^48^ software was used to generate a tree from the multiple sequence alignment using default parameters. The iTOL^49^ software was used for tree visualization. A defense score was calculated for each clade as described above, while removing the effector genes from the positive set to avoid artificial inflation of the scores.

### Bacterial strains, phages and growth conditions

*Escherichia coli* strains (MG1655, DH5α, and Δ*iscR*, Δ*recB*, Δ*recC* and Δ*recD* from the Keio collection^12^) were grown in LB or LB agar at 37°C unless mentioned otherwise. Whenever applicable, media were supplemented with ampicillin (100 μgml^−1^) or kanamycin (50 μgml^−1^) to ensure the maintenance of plasmids.

Phage SECphi4 was isolated from sewage samples as previously described^19^ on *E. coli* MG1655. Phage DNA was extracted using the Qiagen DNeasy blood and tissue kit (cat #69504) and DNA libraries were prepared using a modified Nextera protocol^50^ for Illumina sequencing. Phage DNA was assembled from sequenced reads using using SPAdes v. 3.10.1 (with the -careful and -cov-cutoff auto modifiers)^51^. The assembled genome was deposited in GenBank under the accession number MT331608 (Supplementary Table 6). Classification of the phage family was performed according to the most closely related known phage based on sequence similarity, as described previously^19^.

### Plasmids and strain construction

Primers used in this study are shown in Supplementary Table 3. Plasmids built for this study and the process of their construction are presented in Supplementary Tables 4 and 5. Retron systems (pAA1-pAA22) were synthetized and cloned in plasmid pSG1-RFP (between the AscI and NotI sites of the multiple cloning sites) by Genscript Corp. as previously described^19^ (Supplementary Table 4). Plasmids used for msDNA isolation, pAA23 and pAA56, were constructed through Gibson assemblies that were first transformed into *E. coli* DH5α and then into *E. coli* MG1655 (Supplementary Table 5).

Plasmids pAA63, pAA64, pAA69, pAA73 and pAA74 used for the expression of phage genes in *E. coli*, were constructed through Gibson assemblies presented in Supplementary Table 5. Plasmid pBbS8k-RFP (obtained from Addgene, Plasmid #35276^52^) was used as vector and amplified by primers AS_328, AS_329. All phage genes (Supplementary Table 5, the “Insert” rubrics) were cloned in the same position, under the control of the pBAD inducible promoter. Phages genes were amplified by PCR on phage DNA from WT or mutant phages (Supplementary Table 5) using the primers listed in Supplementary Table 3. After Gibson assembly, these plasmids expressing phage genes were first transformed into *E. coli* DH5α. As some toxicity could be expected, 1% glucose was added to the recovery media and in the selecting plates to avoid leaky expression of potentially toxic genes. Plasmids were then transformed into the relevant strains of *E. coli* MG1655 (with or without Ec48). 1% glucose was added also here to the recovery media and in the selecting plates to avoid leaky expression of potentially toxic genes.

### Plaque assays

Phages were propagated by picking a single phage plaque into a liquid culture of *E. coli* MG1655 grown at 37°C to OD_600_ 0.3 in LB medium supplemented with 0.1 mM MnCl_2_ and 5 mM MgCl_2_ until culture collapse, after which the culture was centrifuged for 10 minutes at 4000 r.p.m and the supernatant was filtered through a 0.2 μM filter to get rid of remaining bacteria and bacterial debris. Lysate titer was determined using the small drop plaque assay method as described in ref^53^.

Plaque assays were performed as previously described^19,53^. Bacteria (*E. coli* MG1655 with plasmids pAA1-pAA22) or negative control (*E. coli* MG1655 with a plasmid pSG1-RFP) were grown overnight at 37°C. Then 300 μl of the bacterial culture was mixed with 30 ml melted MMB agar (LB + 0.1 mM MnCl2 + 5 mM MgCl_2_ + 0.5% agar) and let to dry for 1 hour at room temperature. 10-fold serial dilutions in MMB was performed for each of the 12 tested phages and 10 μl drops were put on the bacterial layer. Plates were incubated overnight at 25°C (for phages SECphi4, SECphi6, SECphi18, SECphi27, SECphi17, and T7) or 37°C (for phages λ-vir, P1, T2, T4, T5, and T6). Efficiency of plating (EOP) was measured and compared between the defense and control strain.

### Detection of non-coding RNA expression using RNA-seq

*E. coli* MG1655 cells containing plasmid pAA1 were diluted 1:100 in 5 ml LB medium supplemented with antibiotics (ampicillin). These cells were grown at 37°C with shaking at 250 r.p.m. to an OD_600_ of 0.6. Samples were centrifuged for 10 minutes at 4000 r.p.m at 4°C. The supernatant was discarded, and pellets were used for RNA extraction.

Bacterial pellets were treated with 100 μl of 2 mg/ml lysozyme using Tris 10 mM EDTA 1 mM pH 8.0 as a buffer. Samples were incubated at 37°C for 5 minutes. 1ml of TRI-reagent (Sigma-Aldrich, 93289) was added to each sample. Samples were then vortexed for 10 seconds to promote lysis before addition of 200μl chloroform. Following another vortexing step, the samples were left at room temperature for 5 minutes to allow phase separation. Samples were then centrifuged at 12,000 g at 4°C for 15 minutes. The upper phase was collected and 500 μl of isopropanol was added. Samples were then incubated overnight at −20°C. The next day, samples were washed. Following 30 minutes of centrifugation at 12000 g at 4°C, the supernatant was discarded leaving a small pellet. 750 μl of ice cold 70% ethanol was added without disturbing the pellet and the samples were then centrifuged for 10 minutes at 12,000 g at 4°C. This wash was repeated once after which the remaining ethanol was discarded and the tubes were left open for 5 minutes to remove residual ethanol. The pellet was then resuspended in 50 μl water and incubated for 10 minutes at 56°C to promote elution. RNA concentrations were measured using Nanodrop.

All RNA samples were treated with TURBO™ DNase (Life technologies, AM2238). Ribosomal RNA depletion and RNA-seq libraries were prepared as described in ref^54^, except that all reaction volumes were reduced by a factor of 4. RNA-seq libraries were sequenced using Illumina NextSeq platform. Reads were mapped as described in^54^ to the reference genome of *E. coli* MG1655 (GenBank accession no. U00096.3) as well as to the sequence of plasmid pAA1 (Supplementary Table 4). For *Paenibacillus polymyxa* ATCC 842, reads were found in the European Nucleotide Database (ENA), study accession no. PRJEB34369, and mapped to the reference genome of *Paenibacillus polymyxa* ATCC 842 (genome ID 2547132099 in the IMG database^25^).

### Isolation of msDNA

msDNA isolation was performed as described in ref^55^. Despite high expression of ncRNA, detection of msDNA from *E. coli* requires the overexpression of the RT and cognate ncRNA. For this, plasmids pAA23 and pAA56 were designed to encode the RT and ncRNA of Retron-Eco8 or retron Ec78, respectively, under the control of an inducible pBAD promoter (pBAD/His A, Thermofisher, Catalog number 43001) (Supplementary Table 5). Retron Ec78 was chosen as a positive control because of the relatively high expression of its msDNA as compared to other *E. coli* retrons^55^. The same plasmid encoding a GFP instead of the retron was used as negative control.

*E. coli* MG1655 cells containing plasmids pAA23, pAA56 or the negative control were diluted 1:100 in 25 ml LB medium supplemented with antibiotics (100 μg/ml ampicillin). After 30 min of growth at 37°C with shaking of 250 r.p.m, expression of the retron was induced with 0.2% arabinose (final concentration) and grown to an OD_600_ of 0.7. Samples were centrifuged for 10 minutes at 4000 r.p.m at 4°C, after which the supernatant was discarded.

To isolate the msDNA, each pellet was treated with three solutions: Solution I (50 mM glucose, 10 mM EDTA and 25 mM Tris-HCl, pH 8.0), Solution II (0.2 N NaOH, 1% SDS) and Solution III (3 M potassium acetate, 2 M acetic acid). 200 μl of Solution I was added to each pellet followed by vortex. Then, 200 μl of Solution II was added and the tubes were mixed by inverting each tube 5 times. 400 μl of Solution III was then added and samples were mixed again by inverting the tubes 5 times. Samples were centrifuged for 10 minutes at 4000 r.p.m at 4°C, after which the supernatant was collected.

Nucleic acids were then precipitated using ethanol. 2.1 ml of 100% ethanol was added to 700 μl of the collected supernatant to allow precipitation. Samples were centrifuged at 4°C, 13,000 r.p.m for 30 minutes and the supernatant was discarded. To wash the pellets, 2.1mL of 70% ethanol was added, without disturbing the pellet, and the samples were centrifuged at 4 °C, 13,000 r.p.m for 10 minutes. Supernatant was discarded and samples were dried for 20 minutes before resuspension in 20 μl water.

An RNAseA treatment was then applied. 2ul of RNAseA 10 μg/μl (from Qiagen Miniprep kit Cat No./ID: 27106) was added to each sample. Samples were incubated at 37°C for 20 minutes. To facilitate visualization on a gel, the ssDNA ladder was diluted 1:100 and the positive control (pAA56, Ec78) was diluted 1:10. 10ul of each of these samples was then mixed with 2X TBE urea loading dye and incubated at 70°C for 3 minutes. After 5 minutes of cooling down, samples were loaded on a 10% denaturing polyacrylamide gel and run for 1h30 minutes at 180V. For visualization, the gel was washed once with TBE, then incubated for 10 minutes in 10% acetic acid, followed by a 20 minutes incubation in an ethidium bromide bath for staining.

### Isolation of mutant phages

To isolate mutant phages that escape Ec48 defense, phages were plated on bacteria expressing the Ec48 defense system (*E. coli* MG1655 with plasmid pAA10, Supplementary Table 4) using the double-layer plaque assay^56^. MMB 1.1% agar was used as the bottom layer, and MMB 0.3% or 0.5% agar was used for the top layer (for λ-Vir and T7, respectively). For the λ-vir phage, the double-layer plates were incubated overnight at 37°C, and 10 single plaques were picked into 90 μl phage buffer (50 mM Tris pH 7.4, 100 mM MgCl2, 10 mM NaCl). For phage T7, the double-layer plate was incubated overnight at room temperature and the entire top layer was scrapped into 2 ml of phage buffer to enrich for phages that escape Ec48 defense. The phages were left for 1 h at room temperature during which the phages were mixed several times by vortex to release them from the agar into the phage buffer, after which the phages were centrifuged at 3200 g for 10 min to get rid of agar and bacterial cells, and the supernatant was transferred to a new tube.

To test the phages for the ability to escape from Ec48 defense, the small drop plaque assay was used^53^. 300 μl bacteria −Ec48 (*E. coli* MG1655 with plasmid pAA10) or negative control (*E. coli* MG1655 with a plasmid pSG1, lacking the Ec48 defense system) were mixed with 30 ml melted MMB 0.3% or 0.5% agar (for λ-Vir and T7, respectively) and let to dry for 1 hour at room temperature. 10-fold serial dilutions in phage buffer was performed for the ancestor phages (used for the original double layer plaque assay) and the phages formed on Ec48 and 10 μl drops were put on the bacterial layer. The plates were incubated overnight at 37°C (for λ-vir phage) or at room temperature (for T7 phage). Efficiency of plating (EOP) was measured and compared between the defense and NC strain.

### Amplification of mutant phages

Isolated phages for which there was decreased defense compared to the ancestor phage were further propagated by picking a single plaque formed on Ec48 in the small drop plaque assay into a liquid culture of Ec48 cells grown in 1 ml MMB to an OD_600_ of 0.3. The phages were incubated with the bacteria at 37°C 200 r.p.m for ~3 hr, and then an additional 9 ml of bacterial culture grown to OD_600_ 0.3 in MMB was added, and incubated for an additional ~3 h at 37°C with shaking at 200 r.p.m. The lysate was then centrifuged at 3200 g for 10 min and the supernatant was filtered through a 0.2 μM filter to get rid of remaining bacteria.

Phage titer was then checked using the small drop plaque assay on the negative control strain and in cases where the titer was less than 10^7^ pfu/ml, the phage titer was raised using either propagation in liquid culture (see plaque assays section above for detailed protocol) or plate lysate^56^ on Ec48 cells. For plate lysate, 100 μl of phage lysate containing 10^3^−10^5^ plaque forming units (pfu) was mixed with 100 μl Ec48 cells and left at room temperature for 10 minutes, after which 5 ml of pre-melted 0.3% MMB was added and then poured onto a bottom layer of MMB. The phages were incubated with the bacteria overnight at 37°C (for λ-vir phage) or at room temperature (for T7 phage), letting the phage to lyse the cells. Then the entire top layer was scrapped into 5 ml of phage buffer, and for 1 h at room temperature during which the phages were mixed several times by vortex to release them from the agar into the phage buffer. The phages were centrifuged at 3200 g for 10 min to get rid of agar and bacterial cells, and the supernatant was filtered through a 0.2 μM filter.

### Sequencing and genome analysis of phage mutants

High titer phage lysates (>10^7^ pfu/ml) of the ancestor and isolated phage mutants were used for DNA extraction. 500 μl of the phage lysate was treated with DNAse-I (Merck cat #11284932001) added to a final concentration of 20 μg/ml and incubated at 37°C for 1 hour to remove bacterial DNA. DNA was extracted using the Qiagen DNeasy blood and tissue kit (cat #69504) starting from the Proteinase-K treatment step to lyse the phages. Libraries were prepared for Illumina sequencing using a modified Nextera protocol as previously described^50^. Reads were aligned to the phage reference genomes (GenBank accession numbers: NC_001416.1 (λ phage), NC_001604.1 (T7 phage)) and mutations compared to the reference genome were identified using Breseq (version 0.29.0) with default parameters^57^. Only mutations that occurred in the isolated mutants, but not in the ancestor phage, were considered. Silent mutations within protein coding regions were disregarded as well.

### Bacterial growth upon induction of phage genes

A single fresh colony was picked into 5 ml of LB supplemented with ampicillin (100 μgml^−1^), kanamycin (50 μgml^−1^) and 1% glucose to avoid leaky expression from the pBAD promoter, and grown at 37°C overnight. Bacteria were diluted 1:50 into 3 ml fresh media (LB with ampicillin, kanamycin and 1% glucose) and grown at 37°C until the bacterial cultures reached an OD_600_ of 0.1.

For the induction of T7 gp5.9 and λ-vir Gam, after reaching an OD_600_ of 0.1, 180 μl of the bacteria were then dispensed into a 96-well plate and incubated at 37°C with shaking in a TECAN Infinite200 plate reader with OD_600_ measurement every 6 minutes. When bacteria reached an OD_600_ of 0.3, 20 μl of ultra-pure water (for uninduced samples) or 3% arabinose (for induced samples, for a final concentration of 0.3%)), was added to the bacterial cultures and the bacteria were then incubated further at 37°C with shaking in a TECAN Infinite200 plate reader with OD_600_ measurement every 6 minutes.

For the induction of T7 gp1.7, after reaching an OD_600_ of 0.1, the bacteria were centrifuged at 3200 g 4°C for 10 min, the supernatant was removed and the bacteria were resuspended in 1 ml fresh LB to wash away remaining glucose (for optimal induction of gp1.7 under the pBAD promoter). The bacteria were then centrifuged again, after which the supernatant was discarded and the bacteria were resuspended in 10% of the initial volume (300 μl) of LB. 20 μl of the concentrated bacteria were then transferred to a 96-well plate containing 180 μl LB supplemented with ultra-pure water (for uninduced samples) or 0.3% arabinose (for induced samples). The plate was then incubated at 37°C with shaking in a TECAN Infinite200 plate reader with OD_600_ measurement every 6 minutes.

### CFU count experiments

For CFU count experiments, bacteria were grown overnight at 37°C in 5 ml of LB supplemented with ampicillin (100 μgml^−1^), kanamycin (50 μgml^−1^) and 1% glucose. Bacteria were diluted 1:50 into fresh media (as above), incubated at 37°C with shaking at 200 r.p.m and OD was monitored every ~30 minutes. When bacteria reached an OD_600_ of 0.3, 3 ml of the 6 ml culture was transferred to a new tube, and 333 μl of either ultra-pure water or 3% arabinose was added to each of the tubes (for uninduced and induced respectively). At several time points throughout the experiment, samples were taken for CFU counts. For this, 10 μl of the bacterial culture was taken into 90 μl LB, and 10 μl of serial dilutions of the bacteria in LB were dropped on LB agar plates supplemented with ampicillin (100 μgml^−1^), kanamycin (50 μgml^−1^) and 1% glucose. The plates were then tilted to allow the drops to spread into a line on the plate, and incubated overnight at 37°C after which single colonies were counted.

### Transformation efficiency assay

Strains from the *E. coli* Keio collection^58^ (Δ*recB*, Δ*recC*, Δ*recD* and Δ*iscR* as a negative control) were made electro-competent as follows: cells were grown until OD_600_ of 0.4. Cells were then washed twice with ice-cold water, then once with 10% cold glycerol, and resuspended in 1/100 of their volume in 10% cold glycerol. 100 ng of plasmid pAA1 (containing Retron-Eco8), pAA10 (Ec48) or pSG1 (control) were electroporated in 30 μL of competent cells. Cells were then incubated in 500 μl LB for 1 h at 37°C and plated on LB plates supplemented with kanamycin (50 μgml−1) and ampicillin (100 μgml−1). Transformation efficiency was assessed by counting single colonies formed after overnight incubation at 37°C.

---

**Supplementary Figure 1.**
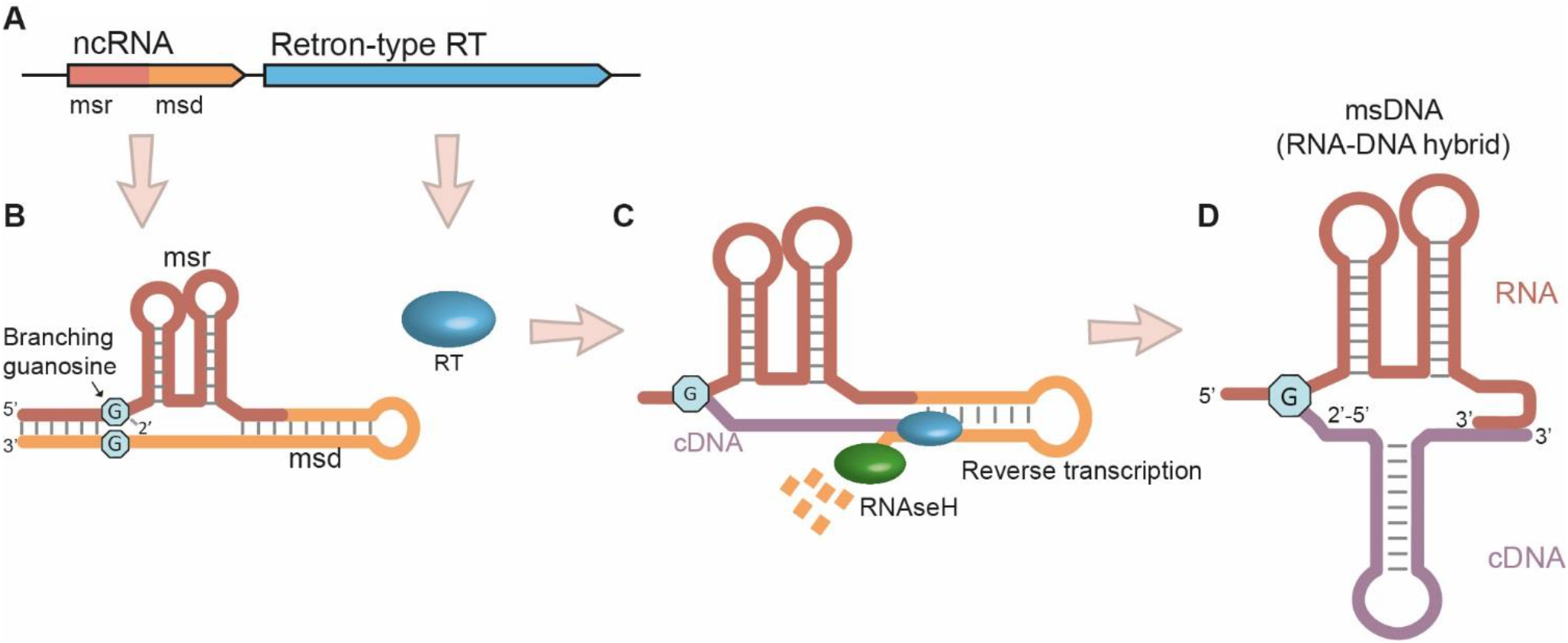
A schematic depiction of retron msDNA synthesis. (A) Retrons encode a non-coding RNA (ncRNA) and a reverse transcriptase (RT). The ncRNA is composed of two parts termed msr and msd. (B) The ncRNA folds such that the edges of the msr and msd base-pair to form a stem that ends with two conserved unpaired guanosine residues on both strands. (C) The RT recognizes the structured ncRNA, uses the 2’OH of the conserved guanosine as primer, and reverse transcribes the msd section which serves as a template, starting from the other conserved guanosine. While the RT forms the cDNA, the template is degraded by RNaseH. (D) This process results in a unique covalently linked RNA-cDNA hybrid called msDNA, which is branched from the guanosine nucleotide. Image was adapted from Darmon et al.^59^.

**Supplementary Figure 2.**
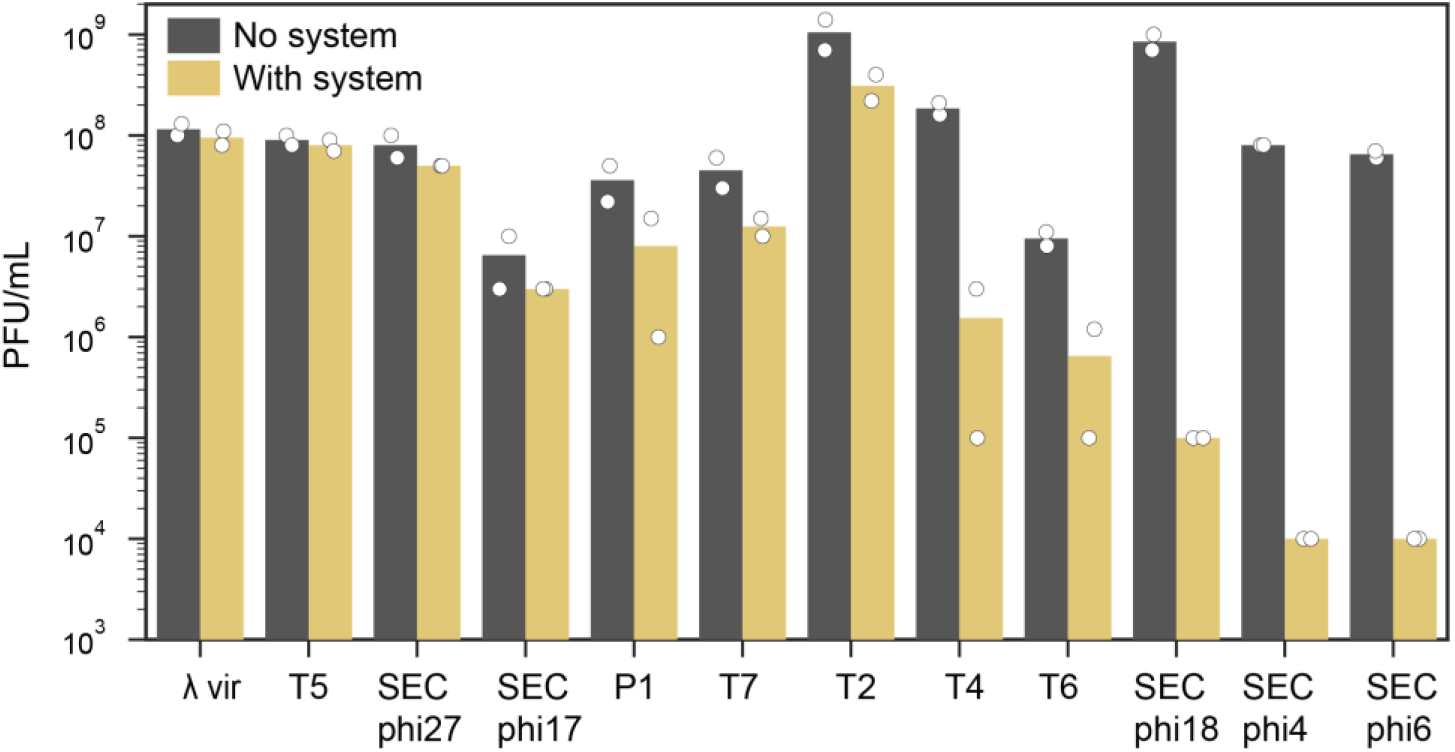
Efficiency of plating of phages infecting *E. coli* with and without the ATPase-RT defense system. The system was cloned from *E. coli* strain 200499, together with its flanking intergenic regions, into the laboratory strain *E. coli* MG1655 (Methods). The efficiency of plating is shown for 12 phages infecting the control *E. coli* MG1655 strain (with a plasmid encoding RFP as negative control) (no system, grey) and the two-gene cassette cloned from *E. coli* 20099 (with system, yellow). Data represent plaque-forming units (PFU) per ml; bar graph represents an average of two replicates, with individual data points overlaid.

**Supplementary Figure 3.**
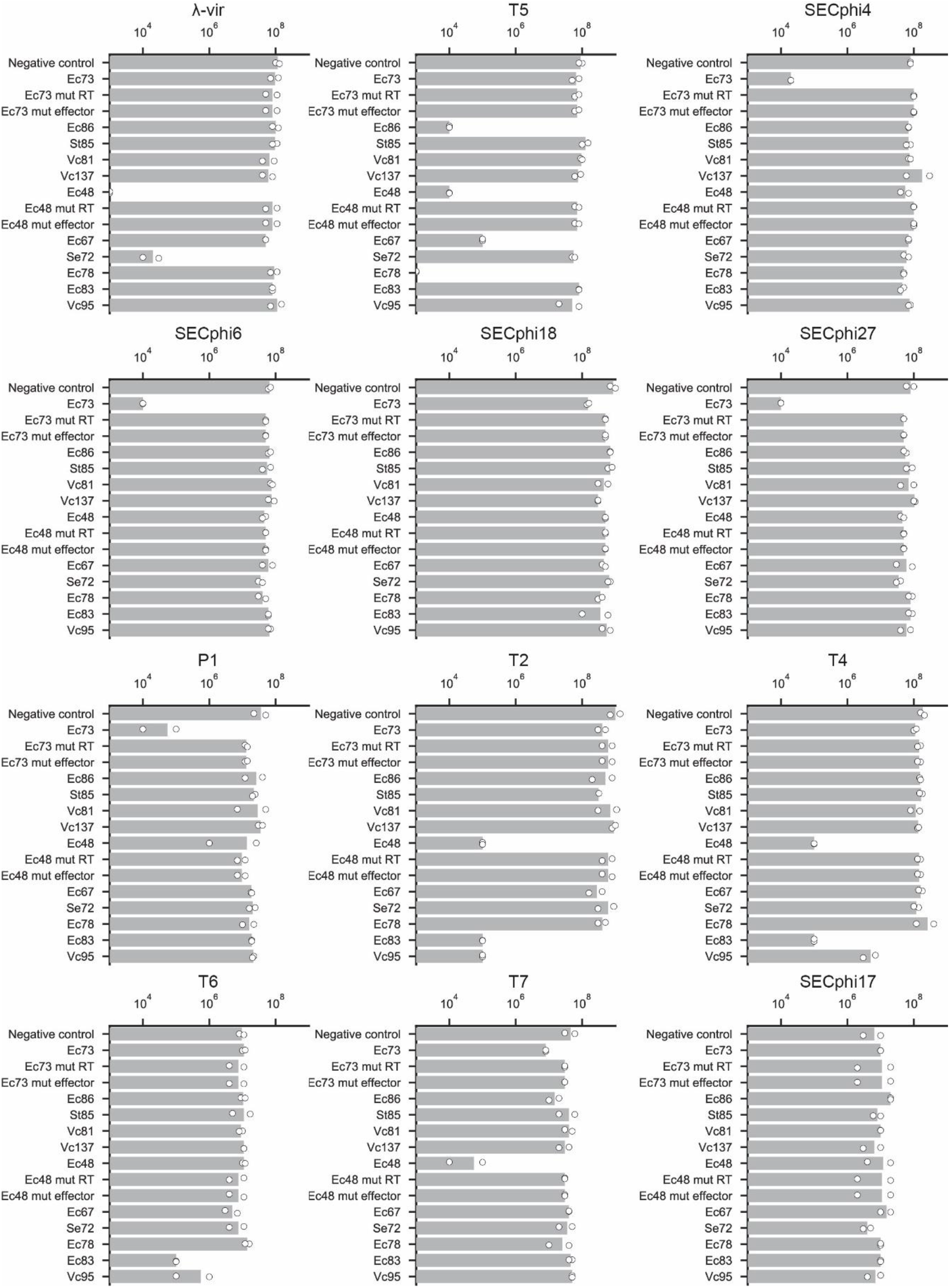
Efficiency of plating of phages infecting retron containing strains. The efficiency of plating is shown for 12 phages infecting *E. coli* MG1655 strains cloned with retron defense systems, or with a plasmid encoding RFP as negative control. Data represent plaque-forming units (PFU) per ml; bar graph represents an average of two replicates, with individual data points overlaid.

**Supplementary Figure 4.**
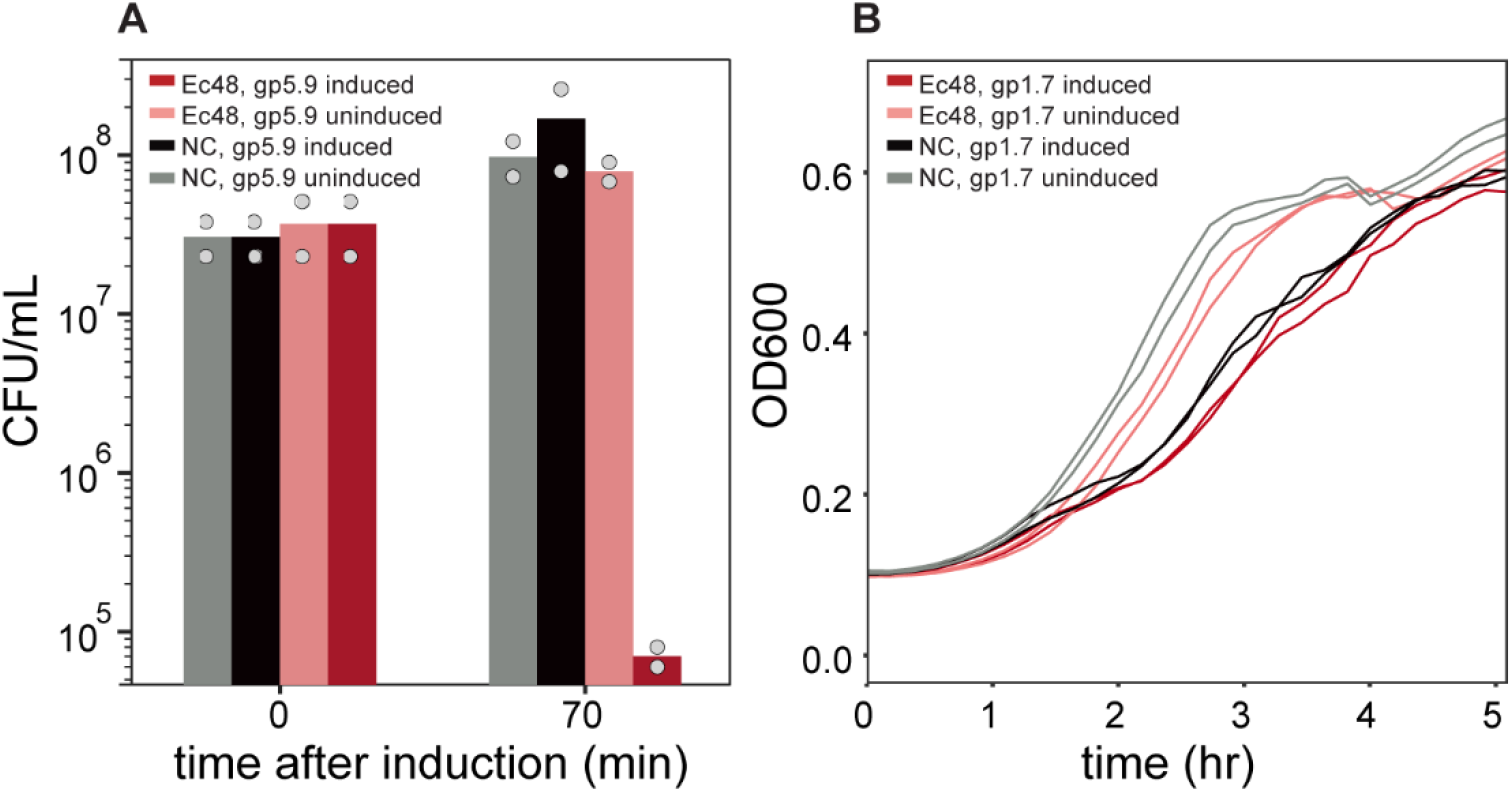
(A) Expression of T7 gp5.9 in cells with the Ec48-containing defense system activates cell death. CFU/ml measurements taken immediately upon (0 min) or 70 minutes after the induction of gp5.9 expression. Induction of gp5.9 was performed by the addition of 0.3% arabinose to exponentially growing cells at OD_600_ 0.1. Ec48, cells containing the Ec48 defense system. NC, negative control cells containing an empty vector instead. Bar graph represents an average of two replicates, with individual data points overlaid. (B) Growth curves of *E. coli* expressing the Ec48 system or a negative control vector that lacks the system (NC), upon induction of phage dNMP kinase (T7 gp1.7 protein) by the addition of 0.3% arabinose at OD_600_ 0.1. Two biological replicates are presented as individual curves.

## Supplementary Tables

**Supplementary Table 1**: RT homologs of known retrons. Attached as an excel file.

---

**Supplementary Table 2:** Mutations observed in phages that escape Ec48 defense. Lambda-vir (NC_001416.1) and T7 (NC_001604.1) genomes were used as references. "*fs*" designates a frameshift beginning at the designated protein residue.

| Phage | Mutant #<br>in figure 4A<br>(where applicable) | Mutated<br>gene name | Gene function | Mutation<br>coordinate<br>in phage<br>genome | Mutation | Mutation effect<br>on protein |
| --- | --- | --- | --- | --- | --- | --- |
| Lambda | 1 | GamS | host RecBCD inhibitor protein | 33006 | $\Delta T$ | *G38fs* |
| | | intergenic<br>region | Rz, cell lysis protein | 46430<br>46433 | T→C<br>C→A | NA |
| | | Bor | Serum Resistance | 46464<br>46668<br>46670 | G→T<br>C→T<br>G→T | Q97K<br>A29T<br>P28Q |
| | | intergenic<br>region | lambdap78, putative envelope<br>protein | 46863<br>46957<br>46985<br>46992<br>47004<br>47669 | T→C<br>+A<br>C→T<br>C→T<br>G→A<br>T→C | NA |
| | | lambdap78 | putative envelope protein | 47143<br>47398<br>47509<br>47529 | C→T<br>C→T<br>T→C<br>C→T | V145I<br>D60N<br>T23A<br>R16K |
| Lambda | 2 | GamS | host RecBCD inhibitor protein | 32947 | $\Delta C$ | *A57fs* |
| Lambda | 3 | GamS | host RecBCD inhibitor protein | 33006 | $\Delta T$ | *G38fs* |
| Lambda | 4 | GamS | host RecBCD inhibitor protein | 32947 | +C | *A57fs* |
| Lambda | 5 | GamS | host RecBCD inhibitor protein | 32947 | $\Delta C$ | *A57fs* |
| Lambda | 6 | GamS | host RecBCD inhibitor protein | 33006 | $\Delta T$ | *G38fs* |
| T7 | 1 | gp1.7 | nucleoside monophosphate kinase | 8548 | A→G | Y128C |
| | | gp5.9 | host RecBCD inhibitor protein | 17426 | T→C | L23P |
| T7 | 2 | gp1.7 | nucleoside monophosphate kinase | 8548 | A→G | Y128C |
| | | gp5.9 | host RecBCD inhibitor protein | 17426 | T→C | L23P |

**Supplementary Table 3:**
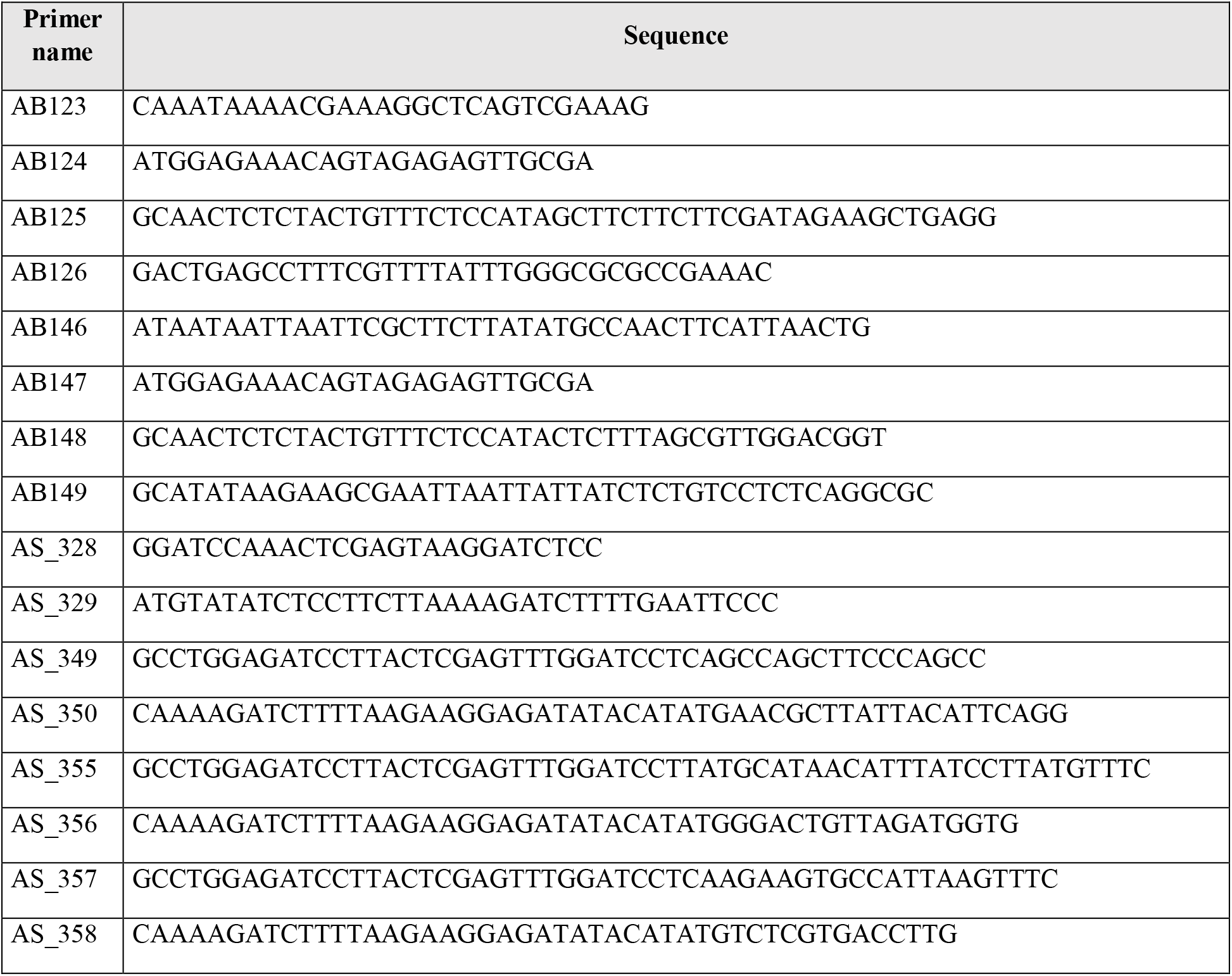
Primers used in this study

**Supplementary Table 4: Sequences synthetized for testing of known retrons**

Attached as an Excel file. Sequences were synthetized and cloned by Genscript Corp, into pSG-RFP1 between the AscI and NotI sites of the multiple cloning site (as described in 19).

**Supplementary Table 5:** Cloning details for plasmids built in this study.

| Plasmids |  | Insert |  | Vector |  |
| --- | --- | --- | --- | --- | --- |
| Name | Description | Primer | Template | Primer | Template |
| pAA23 | pBad-overexpression-new-system | AB125+AB126 | pAA1 | AB123+AB124 | pBad/His A (Thermofisher, Catalog number 43001) |
| pAA56 | pBad-overexpression-Ec78 | AB148+AB149 | pAA13 | AB146+AB147 | pAA23 |
| pAA63 | pBAD-Lambda_Gam | AS_349+AS_350 | Lambda-vir WT | AS_328+AS_329 | pBbS8k-RFP (Addgene, Catalog number 35276) |
| pAA64 | pBAD-Lambda_Gam-A57fs | AS_349+AS_350 | Lambda-vir escapee #4 from Ec48 | AS_328+AS_329 | pBbS8k-RFP (Addgene, Catalog number 35276) |
| pAA69 | pBAD-T7_gp1-7 | AS_355+AS_356 | T7 WT | AS_328+AS_329 | pBbS8k-RFP (Addgene, Catalog number 35276) |
| pAA73 | pBAD-T7_gp5-9 | AS_357+AS_358 | T7 WT | AS_328+AS_329 | pBbS8k-RFP (Addgene, Catalog number 35276) |
| pAA74 | pBAD-T7_gp5-9-L23P | AS_357+AS_358 | T7 escapee #2 from Ec48 | AS_328+AS_329 | pBbS8k-RFP (Addgene, Catalog number 35276) |

**Supplementary Table 6:** Phages used in this study

| Phage | Taxonomy | Accession number |
| --- | --- | --- |
| Lambda-vir | *Siphoviridae* | NC_001416.1 |
| SecPhi17 | *Microviridae* | LT960607.1 |
| SECphi18 | *Siphoviridae* | LT960609.1 |
| SECphi27 | *Siphoviridae* | LT961732.1 |
| SECphi6 | *Siphoviridae* | GCA_902807315 |
| SECphi4 | *Siphoviridae* | MT331608 |
| P1 | *Myoviridae* | AF234172.1 |
| T2 | *Myoviridae* | LC348380.1 |
| T4 | *Myoviridae* | AF158101.6 |
| T5 | *Siphoviridae* | AY543070.1 |
| T6 | *Myoviridae* | MH550421.1 |
| T7 | *Podoviridae* | NC_001604.1 |

## Notes

### Competing Interest Statement

R.S. is a scientific cofounder and advisor of BiomX, Pantheon Bioscience and Ecophage.

